# PlantMDCS: A code-free, modular toolkit for rapid deployment of plant multi-omics databases

**DOI:** 10.64898/2026.02.09.704752

**Authors:** Chen Chen, Yuanyuan Liu, Lei Wang, Jingyi Sai, Yuetian Wang, Wen Yue, Jun Sun, Zixiang Li, Faguo Wang, Jia Tian, Dong Xu, Yuhan Fang

## Abstract

With the rapid accumulation of diverse omics datasets, achieving efficient management and integrative analysis of plant multi-omics data remains a major challenge. Conventional solutions rely on constructing web-based databases, which often demand substantial programming expertise and long-term financial support. To address these limitations, we developed the Plant Multi-omics Database Construction System (PlantMDCS)-a locally deployable, user-friendly, and collaborative platform that unifies database construction and downstream multi-omics analysis within a graphical environment. PlantMDCS adopts a decoupled front-end/back-end architecture. The back end serves as the core engine for data management and computation, and is responsible for the storage, preprocessing, integration, and hierarchical association of multi-omics data. Once initialized, the front end supports the complete research workflow, including data import, querying, integrative analysis and visualization. All operations can be performed without programming, while local resource usage is dominated by disk storage required for user-provided datasets rather than sustained computational overhead. Benchmarking across plant species ranging from *Arabidopsis* to hexaploid wheat demonstrated that database construction can be completed within minutes, independent of genome size or data complexity. PlantMDCS is designed for local deployment to ensure data security, while allowing multi-user collaboration within local networks and supporting controlled remote access for teams distributed across different regions. Overall, PlantMDCS offers a secure and sustainable framework that integrates data management and analysis within a unified system. This design shifts multi-omics research away from fragmented file-based processing toward persistent, database-driven exploration, thereby enhancing analytical efficiency and reproducibility.

## Introduction

In recent years, the rapid advancement of sequencing technologies has driven an exponential growth of multi-omics datasets, including genomic and transcriptomic profiles, population resequencing data, pangenomic resources, metabolomic measurements, and phenotypic traits (Shen et al., 2023; Song et al., 2023a; Song et al., 2023b; Xie et al., 2024; Zavafer et al., 2023). The integrated application of multi-omics data has substantially advanced the identification of the genetic basis underlying key quality traits and has deepened our understanding of evolutionary processes at the genomic level, thereby greatly promoting progress in both botany and agronomy. Such studies often require the simultaneous storage, management, sharing, and analysis of datasets comprising multiple samples, making the effective and efficient decoding of underlying biological infromation a significant challenge for wet-lab biologists and even trained bioinformaticians (Yang et al., 2026).

Currently, the most common approach to this challenge is to establish online public databases or server-hosted repositories for centralized data management, sharing, and routine analysis (Li et al., 2024; Wang et al., 2026). However, this conventional database construction strategy presents several major limitations. First of all, the construction of database systems, such as user interfaces and backend server infrastructures, typically relies on specialized database software. In contrast, data analytical models are usually implemented as scripts written in disparate programming languages. As a result, when handling large-scale datasets for database construction, wet-lab biologists and even bioinformaticians are often required to acquire extensive computational expertise, encompassing not only proficiency in command-line environments but also familiarity with professional database software. Second, depositing sensitive or unpublished datasets on public platforms or cloud services may increase the risk of data leakage or re-identification attacks. This concern is particularly pronounced for genomic data associated with proprietary traits or unpublished plant varieties and species. Third, such systems are often difficult for individual research groups to sustain over the long term, owing to continuous demands of system maintenance, security updates, and financial costs. Therefore, the lack of easily accessible tools for database construction remains a critical bottleneck, and the development of locally deployable, low–technical-barrier, and self-sustaining database platforms has become essential for balancing data security, accessibility, and long-term usability in multi-omics research.

Frameworks such as e!DAL, Flask (Li et al., 2026), Django (Lai et al., 2025), Tripal (Chen et al., 2025), and ThinkPHP (Yang et al.,2023) have attempted to facilitate database construction in the life sciences, among which e!DAL(Arend et al., 2014) and Tripal (Staton et al., 2021) are tailored for research data and plant genome. e!DAL is a lightweight, open-source framework, primarily designed for data storage, sharing, and publication. It integrates key functionalities such as metadata annotation, HTTP(S) access, and WebDAV-based file system support, allowing both local embedded deployment and client–server architecture. However, it does not include advanced data analysis modules commonly found in online biological databases (such as sequence alignment, statistical modeling, or comparative genomics) (Li et al., 2024; Dai et al., 2025), and therefore relies on external plugins or platforms (e.g., VANTED, Galaxy) for analytical capabilities. Tripal, in contrast, was built on the Drupal content management system (CMS) and the Chado database schema, providing a modular and extensible environment for constructing biological databases. Nevertheless, Tripal also lacks integrated analytical functionality for in-depth data exploration, and its core development often depends on short-term research funding, raising concerns about long-term sustainability. Meanwhile, although Tripal supports multiple database backends, its distributed storage and cloud deployment optimization remain underdeveloped, leading to limited scalability and efficiency when handling large-scale datasets compared to systems specifically optimized for high-performance data storage architectures. Moreover, like other frameworks, the steep learning curve of the Drupal framework poses challenges for users without technical backgrounds, making it less accessible to smaller or non-computational research teams.

Here, we present PlantMDCS (Plant Multi-omics Database Construction System), a novel platform designed for the construction and management of locally deployable plant multi-omics databases. Sharply contrast with existing protocols, PlantMDCS adopts a dual-module architecture. The administrator module enables centralized uploading, integration, and management of diverse omics datasets without reliance on external cloud services or public servers, thereby substantially reducing the risk of data leakage. Once data are uploaded, integration is performed automatically, allowing seamless database construction, maintenance, and expansion. The user module provides comprehensive, integrative analysis of the curated datasets and supports the generation of publication-ready, interactive visualizations. All analytical procedures are executed through intuitive graphical operations, such as clicking and dragging, eliminating the need for scripting or advanced computational expertise. Databases constructed with PlantMDCS can be accessed by multiple users through standard web browsers in either local or remote environments, achieving an effective balance between data security, usability, and collaborative accessibility. As such, PlantMDCS is well suited for plant research groups operating with standard computational resources. The development of PlantMDCS was motivated by the need to provide plant researchers with a constructible, shareable, and secure platform for the management and integrative analysis of multi-omics data, enabling efficient data reuse and collaborative exploration while maintaining full control over locally stored datasets.

## Results

### An overview of PlantMDCS

To address the increasing challenges associated with managing, integrating, and securely sharing large-scale plant multi-omics datasets, we developed PlantMDCS (Plant Multi-omics Database Construction System), a locally deployable and collaborative platform that provides functionality comparable to that of established web-based databases, while unifying database construction and downstream multi-omics analysis within a single graphical environment (Figure 1A). PlantMDCS is specifically tailored for plant research groups that require flexible data control, efficient data reuse, and integrative analytical capabilities without dependence on external cloud services or public servers. PlantMDCS is architecturally organized into two complementary modules: an administrator module and a user module, which together support the full life cycle of multi-omics data, from initial integration to biological interpretation. The administrator module is responsible for centralized data uploading, organization, and maintenance, enabling the construction of a standardized and internally consistent multi-omics database (Figure 1B). All datasets are stored and managed locally, ensuring full ownership of raw data and substantially reducing the risk of unintended data exposure. Once data are uploaded, their integration into the database is performed automatically, allowing seamless database initialization, expansion, and long-term maintenance without repeated restructuring. The user module, comprising 46 functions spanning sequence analysis to visualization, is designed to facilitate intuitive access to the curated datasets and supports comprehensive, integrative analyses across multiple omics layers (Figure 1C). Users can explore genomic, transcriptomic, population genomic, metabolomic, and phenotypic data through a unified interface and generate publication-ready visualizations without manual file preparation or scripting. All analytical procedures are executed through simple graphical interactions, such as mouse selection and parameter-free task execution, thereby lowering the technical barrier for researchers with diverse computational backgrounds. Databases constructed using PlantMDCS can be accessed simultaneously by multiple users through standard web browsers in local or remote environments, enabling efficient collaboration within research teams while maintaining strict control over data accessibility. By balancing local deployment, ease of use, and collaborative functionality, PlantMDCS provides a practical and sustainable solution for plant multi-omics data management and analysis, making it suitable for a wide range of plant research applications using standard computational resources.

**Figure 1.**
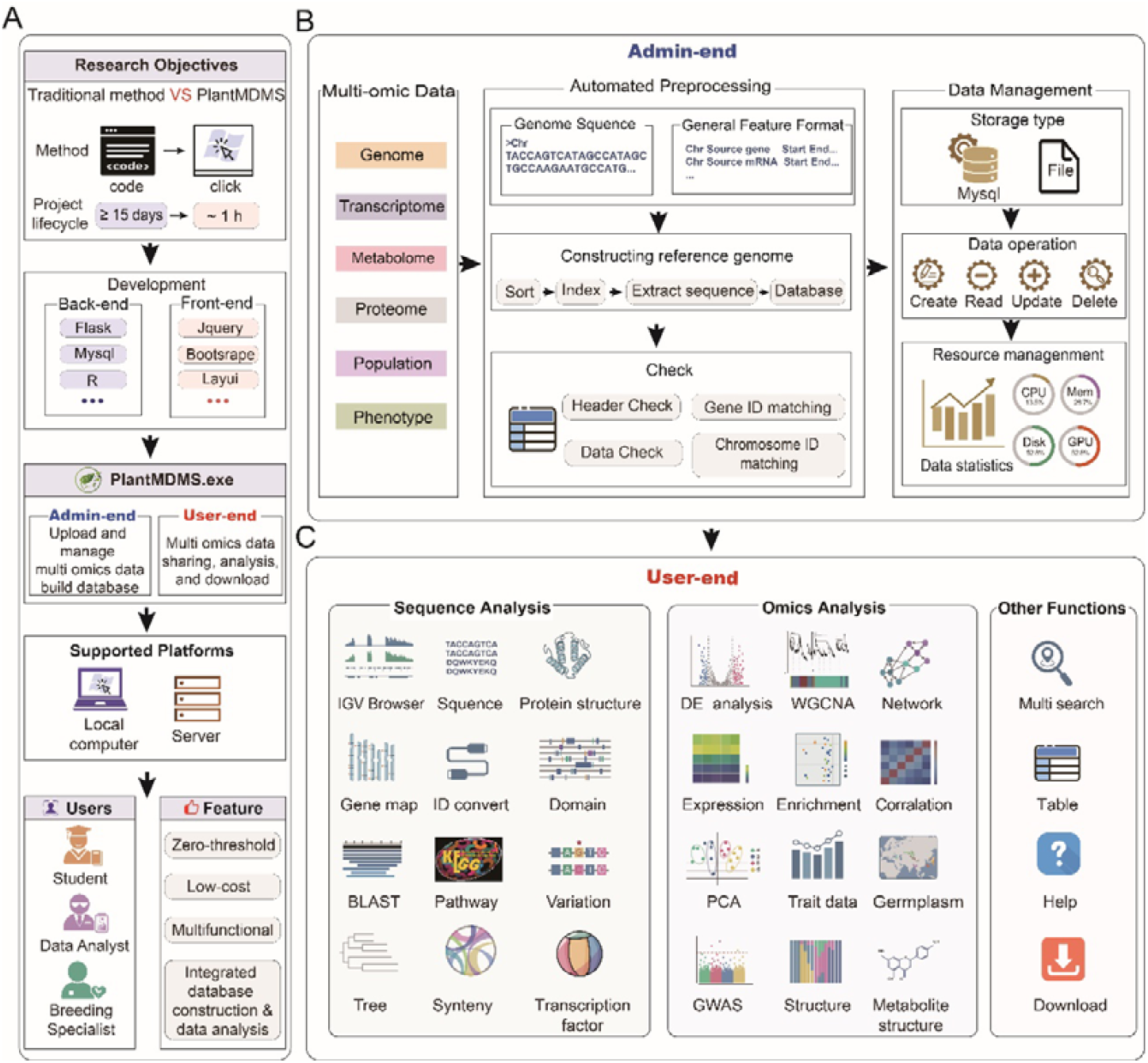
Overview of the PlantMDCS workflow and functional architecture. **(A)** Schematic overview of the core workflow of PlantMDCS, illustrating the integration of data construction, management, and downstream multi-omics analysis within a unified platform. **(B)** Functional layout of the administrator module (Admin-end), which supports data uploading, organization, and centralized management during database construction and maintenance. **(C)** Functional layout of the user module (User-end), enabling integrative exploration and analysis of curated multi-omics datasets through an intuitive graphical interface.

### Design principles of PlantMDCS

PlantMDCS is designed around a genome-centered organizational logic that enables robust and reusable integration of heterogeneous plant multi-omics datasets. The input genome serves as the central anchor of the system, with standardized gene identifiers derived from genome annotation files providing a common reference across data types. Through this structure, genomic features establish logical links among genome sequences, transcriptomic profiles, and population-scale variation data in a manner consistent with the biological flow of genetic information from sequence to expression and variation. Proteomic, metabolomic, and phenotypic datasets are supported as complementary layers that can be managed independently while remaining conceptually aligned with the genome-centered framework (Figure 2A). This strategy is inherently project-independent and species-agnostic, allowing PlantMDCS to accommodate any plant species with an available reference genome without modifying the underlying data organization.

**Figure 2.**
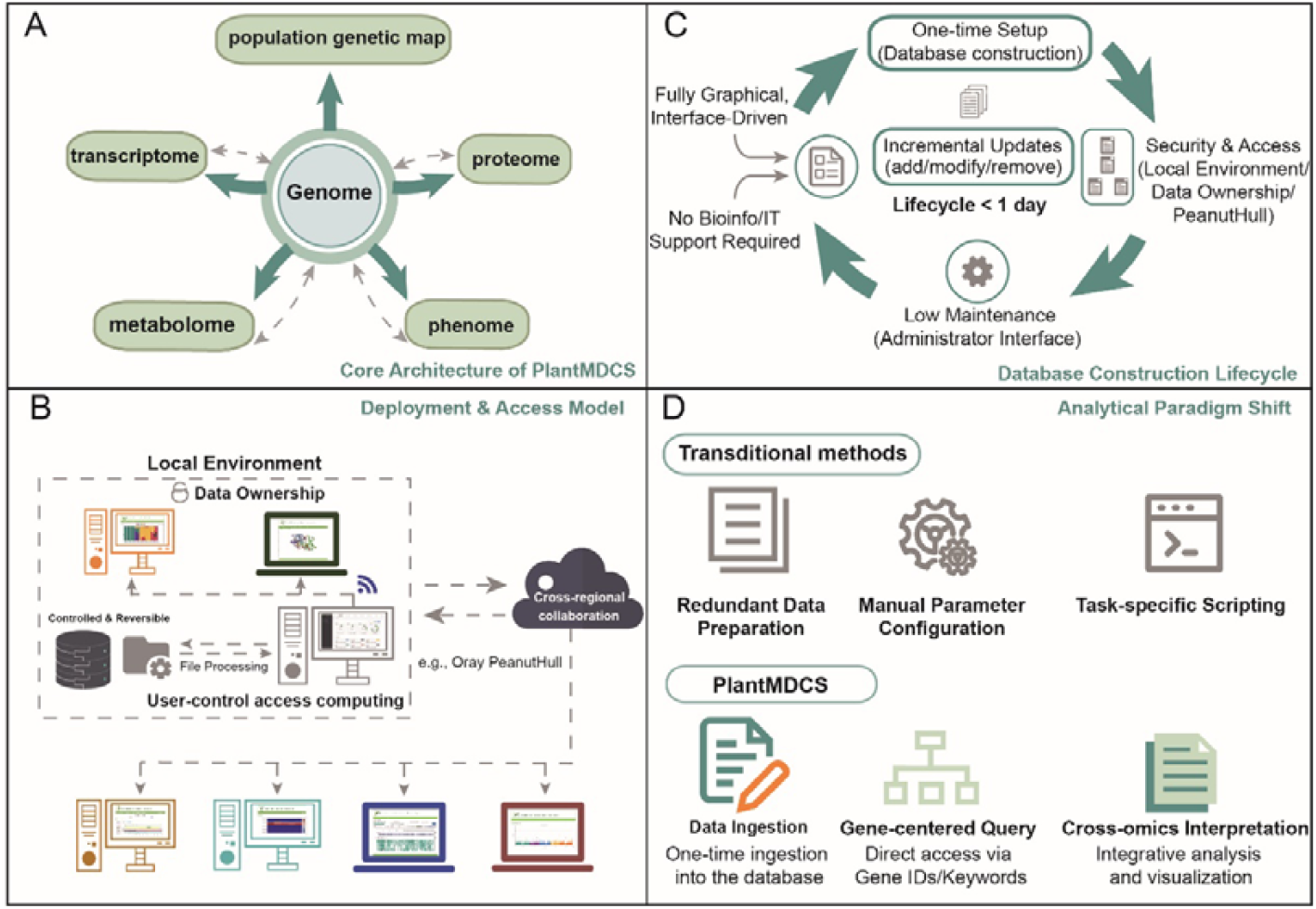
Overview of PlantMDCS architecture and workflow. **(A)** Core architecture: PlantMDCS adopts a genome-centered framework that integrates heterogeneous plant multi-omics datasets through standardized gene identifiers, enabling consistent links among genome, transcriptome, proteome, metabolome, and population variation data. **(B)** Deployment and access: Databases are locally deployed under full user control, ensuring data ownership and security. Controlled external access (e.g., Oray PeanutHull) supports collaborative use without compromising administrative control. **(C)** Database lifecycle: Database construction is a one-time, interface-driven process supporting incremental updates without bioinformatics or IT support. **(D)** Analytical paradigm: Transition from file-based processing to gene-centered, data-driven exploration enables direct querying and cross-omics interpretation within an integrated environment.

To ensure long-term usability and sustainability, PlantMDCS separates database deployment from subsequent data utilization and maintenance through a fully graphical, interface-driven design. Database construction is performed once and can be reused continuously, while newly generated datasets can be incrementally added, modified, or removed without disrupting existing data relationships or requiring structural reconstruction (Figure 2B). All routine management tasks are handled through the administrator interface, eliminating the need for continuous bioinformatics or IT support and ensuring that database maintenance remains feasible despite project expansion or personnel turnover. Storage requirements scale directly with the size of user-provided datasets, whereas computational resources are primarily consumed during user-initiated analyses, with resource usage determined by the analytical tasks selected rather than by background system operations. As a result, PlantMDCS remains operable on standard computational infrastructure commonly available in plant research laboratories.

By emphasizing local deployment combined with flexible access control, PlantMDCS reconciles data security with collaborative accessibility. Databases are constructed and hosted within user-controlled computing environments, ensuring full ownership of both raw and processed data. Within local network environments, multiple users can simultaneously access the database through standard web browsers, enabling real-time internal collaboration while preventing unintended external exposure of sensitive or unpublished datasets. When broader collaboration or data dissemination is required, controlled external access can be enabled in a deliberate and reversible manner (such as, Oray PeanutHull), allowing researchers distributed across different regions to interact with the database without compromising data security or administrative control (Figure 2C).

Rather than functioning as a collection of loosely assembled analytical tools, PlantMDCS introduces a gene-centered and data-driven exploration paradigm that fundamentally alters the analytical workflow. Tranditional file-oriented analysis requires repeated data preparation and parameter configuration for individual tasks, whereas PlantMDCS shifts the analytical focus to integrated biological entities once data have been ingested into the database (Figure 2D). This design allows users to move seamlessly from data querying to integrative analysis and interpretation without redundant file handling. By transitioning from file-based processing to gene-centered multi-omics exploration, PlantMDCS supports more efficient hypothesis generation and cross-omics interpretation, representing a qualitative advance over conventional GUI-based analysis workflows.

### Application-oriented database construction reduces barriers and costs

Existing strategies for constructing plant omics databases are predominantly based on general-purpose web frameworks or server-oriented platforms, which typically require substantial programming expertise, prolonged development cycles, and continuous technical maintenance.

Regardless of the specific framework employed, these approaches share several inherent characteristics: database construction is time-consuming, programming is indispensable for both deployment and customization, and long-term operation depends on sustained bioinformatics or IT support. As a consequence, the overall maintenance cost remains high, and database sustainability is often vulnerable to personnel turnover or limited technical resources (Table 1).

**Table 1.** Comparison of database construction strategies highlighting application barrier and maintenance cost.

| **Evaluation**<br>**Dimension** | **PlantMDS** | **ThinkPHP** | **Tripal** | **Flask** | **Django** |
| --- | --- | --- | --- | --- | --- |
| Construction<br>cycle | ~20 min | > 15 days | > 15 days | > 15 days | > 15 days |
| Programming<br>required | No | Yes | Yes | Yes | Yes |
| Maintenance<br>cost | Zero | High | High | High | High |
| operating<br>platform | Windows | Linux | Linux | Linux | Linux |

Note: time statistics were calculated using a maize multi-omics dataset comprising five genomes (B73, HZS, MEXICANA, Mo17 and SK), 23 transcriptomes, 339 metabolomic datasets, 507 resequencing datasets, and 436 phenotypic datasets, with a total data volume of 9.45 Gb.

In contrast, PlantMDCS adopts a fundamentally different application-oriented strategy that directly targets these bottlenecks. While conventional frameworks emphasize flexibility at the cost of complexity, PlantMDCS prioritizes usability and reusability by providing a fully graphical, zero-code workflow for database construction, management, and routine operation. As reflected in Table 1, tasks that conventionally require manual coding and environment configuration can be completed in PlantMDCS through interface-driven interactions, substantially shortening the database construction cycle and eliminating dependence on programming expertise. In addition, the direct ingestion of large-scale datasets from local files further improves efficiency by avoiding intermediate data transfer and format conversion steps. Based on empirical time statistics, the overall database construction efficiency of PlantMDCS exceeds that of conventional frameworks by more than three orders of magnitude.

This difference in design philosophy also translates into markedly different maintenance costs. In framework-based solutions, routine updates, data expansion, or functional adjustments frequently necessitate technical intervention, resulting in persistently high maintenance overhead. By contrast, PlantMDCS supports incremental data updates and routine database management entirely through the administrator interface, enabling one-time deployment with long-term reuse. As a result, ongoing maintenance does not require dedicated technical personnel, effectively reducing long-term maintenance cost to a negligible level from the user’s perspective.

Finally, the operational environment further distinguishes PlantMDCS from conventional database construction strategies. Whereas most existing solutions are designed for Linux-based server infrastructures, PlantMDCS is deployable on widely available desktop operating systems, lowering both hardware and administrative barriers. This design choice enables plant research groups to establish functional multi-omics databases using standard computational resources, while retaining capabilities comparable to those of established web-based databases. Collectively, these differences demonstrate that the advantage of PlantMDCS lies not in the accumulation of additional functions, but in a substantial reduction of application barriers and long-term cost, achieved through a deliberate shift from code-centric development to user-centric deployment.

### Demonstration of scalability and broad applicability of PlantMDCS across plant species

In the preceding sections, we have outlined the overall functionality of PlantMDCS, described the underlying organizational logic of its design, and compared PlantMDCS with existing tools addressing similar tasks. In this section, we present case studies based on real plant datasets to illustrate the basic operation of PlantMDCS (primarily using *A*.*thaliana* data), and to evaluate whether its performance is affected by differences in plant genome size and multi-omics data complexity.

### The practical workflow of database construction in PlantMDCS

The practical workflow of database construction in PlantMDCS was shown in Figure 3. The process begins with launching the graphical startup interface of PlantMDCS, where essential system parameters, including the service port, administrator credentials, and session identifier, are configured to initialize the local database service (Figure 3A). Once the system is running, administrators access the management environment through the admin-end login interface using authorized credentials (Figure 3B), ensuring controlled access to database construction and maintenance functions. After successful authentication, the database and queue are first configured (Figure 3C). Subsequently, the multi-omics database is initialized by uploading a reference genome sequence together with its corresponding genome annotation file in GFF format (Figure 3D). Furthermore, data standardization can be performed using SPDEv3.0 and GFAP to ensure compatibility with downstream analyses (Xu et al., 2023; Xu et al., 2025). This step establishes the genome-centered framework that serves as the organizational anchor for subsequent data integration. Based on this reference structure, heterogeneous omics datasets can be incrementally incorporated into the database. As an example, transcriptomic expression data are uploaded through standardized interface-driven workflows, illustrating how omics datasets are ingested, parsed, and linked to genome annotations without manual file manipulation or scripting (Figure 3E). All data processing and integration steps are executed automatically in the background, enabling seamless database construction and expansion. Upon completion of data ingestion and integration, the constructed database becomes immediately accessible through the user-end interface (Figure 3F). User-end can browse, query, and analyze integrated multi-omics data through a unified web interface, enabling downstream biological exploration without additional configuration. Together, this workflow demonstrates how PlantMDCS supports the complete life cycle of plant multi-omics database construction (from system initialization and data integration to user-facing analysis) through a fully graphical, locally deployable framework that balances ease of use, data security, and collaborative accessibility.

**Figure 3.**
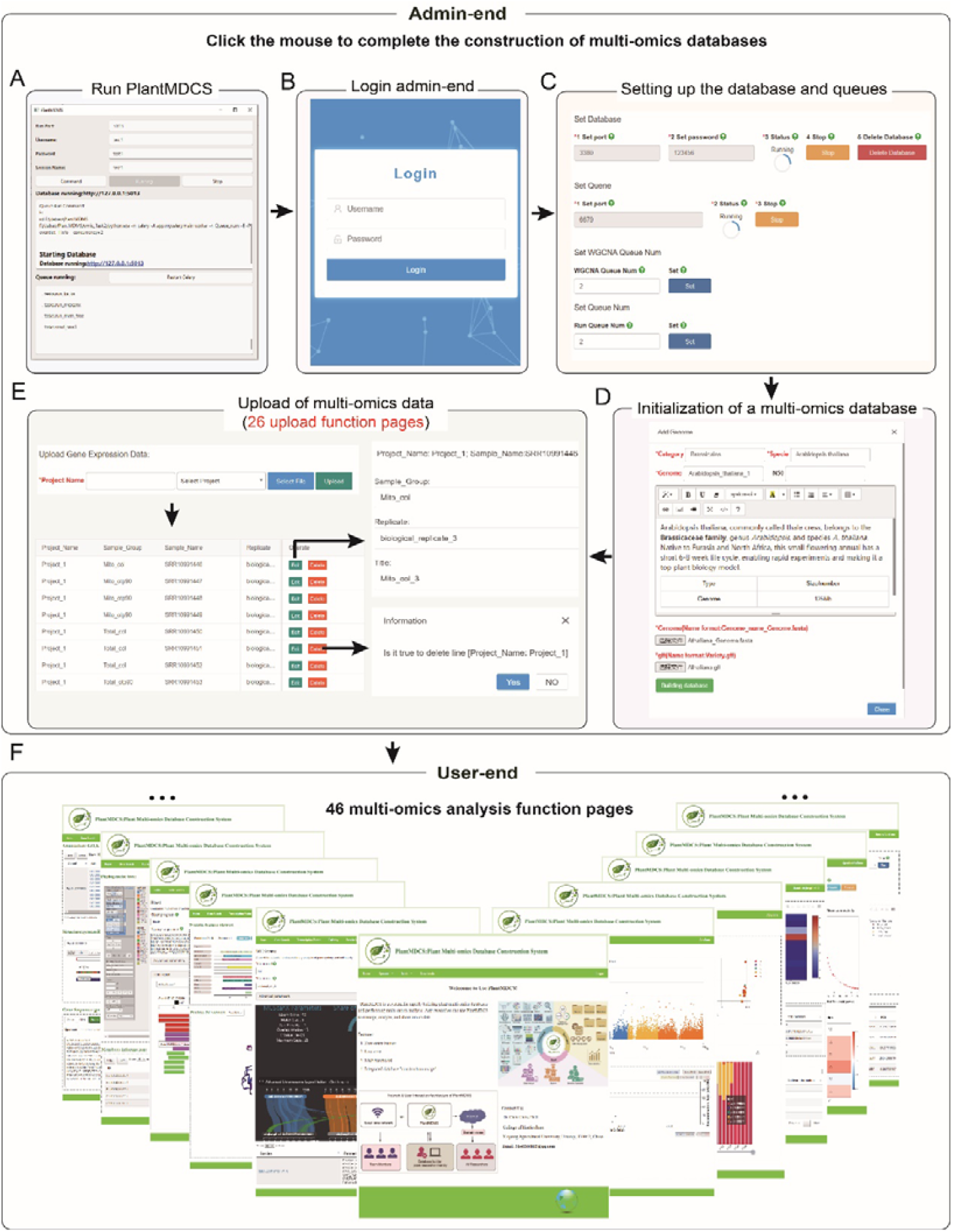
Workflow for constructing a plant multi-omics database using the PlantMDCS admin module. **(A)** Graphical startup interface of PlantMDCS. Users initiate the system by configuring database-related parameters, including the service port, administrator username, login password, and session name. **(B)** Admin-end login interface. Authorized administrators access the management environment by entering valid credentials. **(C)** Configuration of the database environment and task queues at the admin end, enabling subsequent data ingestion and analysis scheduling. **(D)** Initialization of a plant multi-omics database through the upload of reference genome sequence files and corresponding GFF annotation files, establishing the genome-centered framework for downstream data integration. **(E)** Example of multi-omics data ingestion at the admin end, illustrated by uploading transcriptomic expression data, demonstrating the standardized workflow for incorporating heterogeneous omics datasets. **(F)** User-end interface providing access to integrated multi-omics data and downstream analysis functions after database construction. The admin module supports 26 data upload function pages, while the user module provides 46 multi-omics analysis and visualization pages.

### Development and application of function of usr-end in PlantMDCS

The user-end provides 46 functional pages dedicated to multi-omics analyses (Figure 3F). In comparison with conventional analytical approaches, PlantMDCS not only drives the barrier to multi-omics analysis down to zero but also further reduces operational complexity. Taking collinearity analysis as an example (Figure 4), performing collinearity analysis between two species using the One Step MCScanX function in Tbtools requires six sequential steps: 1) inputting the genome sequence of Species A; 2) inputting the GFF file of Species A’s genome; 3) inputting the genome sequence of Species B; 4) inputting the GFF file of Species B’s genome; 5) configuring relevant parameters; and 6) specifying the output path. In contrast, PlantMDCS enables the completion of interspecific collinearity analysis in merely three steps: 1) selecting Genome A; 2) selecting Genome B; and 3) setting the required parameters. Beyond this, PlantMDCS equips users with interactive visualization and in-situ retrieval capabilities. Taking collinearity analysis as a representative example (Figure 4), users can generate an interactive volcano plot upon conducting gene differential expression analysis. By clicking on a data point in the volcano plot, users can initiate an in-system retrieval of the corresponding gene’s annotation information, which is then dynamically displayed on the right side of the volcano plot in real time.

**Figure 4.**
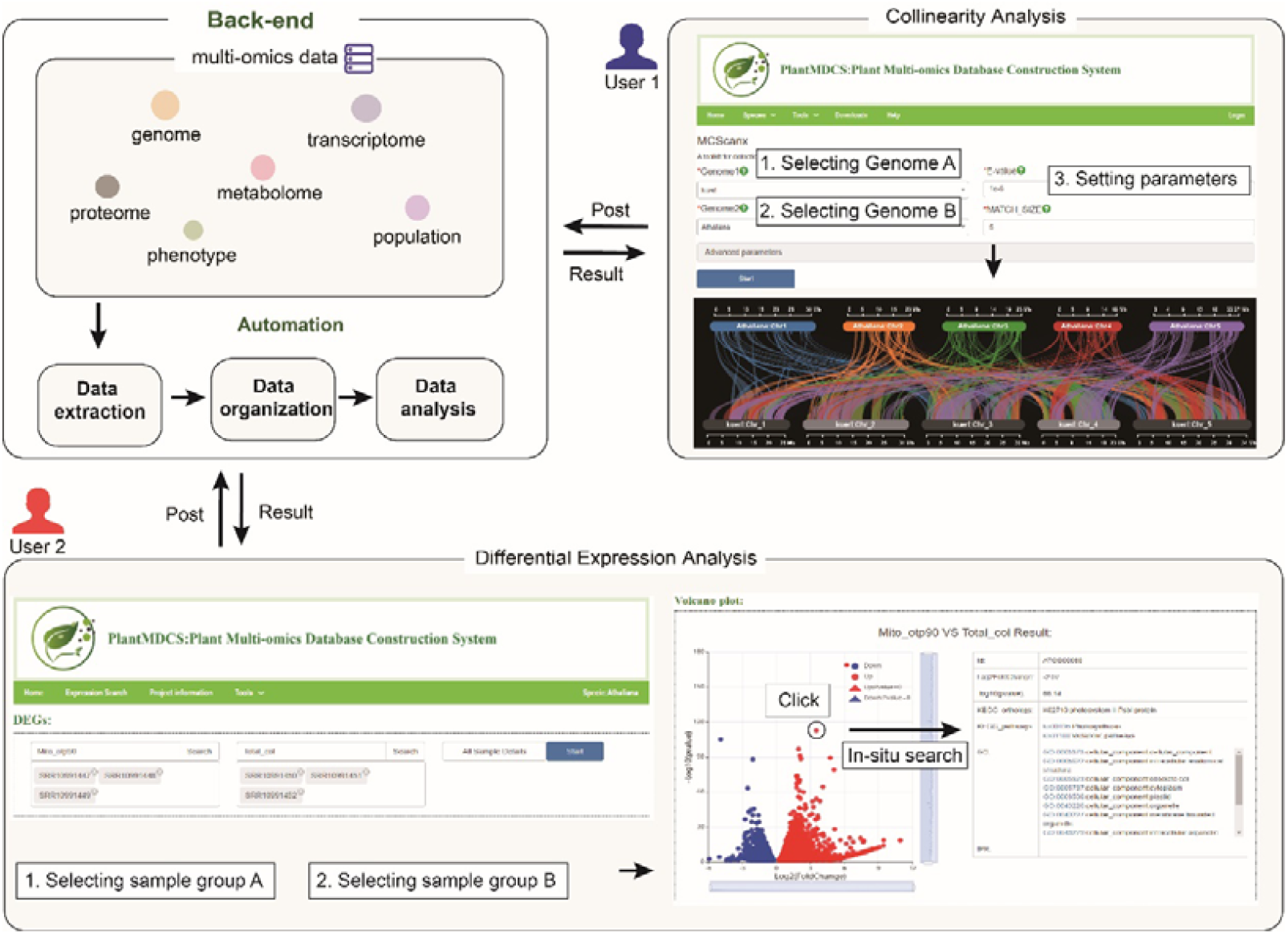
Multi-user analysis and simplified page operation design for PlantMDCS user-end. PlantMDCS allows multi-user access to its user-end via local browsers for multi-omics analyses through POST requests; the system backend automatically performs data extraction, organization and analysis, and returns the results to users.

In addition to the aforementioned two functions, the user-end equips users with a comprehensive set of analytical functions, including sequence retrieval, annotation query, structure visualization (such as protein 3D structure and molecular formula display), expression level inquiry, Weighted Gene Co-expression Network Analysis (WGCNA), genome browser visualization, GO and KEGG enrichment analysis, phenotype query, population structure analysis, Electronic Fluorescent Pictograph (eFP) Browser, heatmap generation, Blast sequence alignment, Principal Component Analysis (PCA), phylogenetic tree construction, gene mapping, collinearity analysis, Manhattan Plot visualization, LD heatmap generation, spatial distribution mapping, transcriptome sequencing visualization, and variant site visualization, etc. (Figure 5). Each function is designed with an extremely user-friendly operational workflow, enabling all users to conduct multi-omics analyses efficiently and effortlessly.

**Figure 5.**
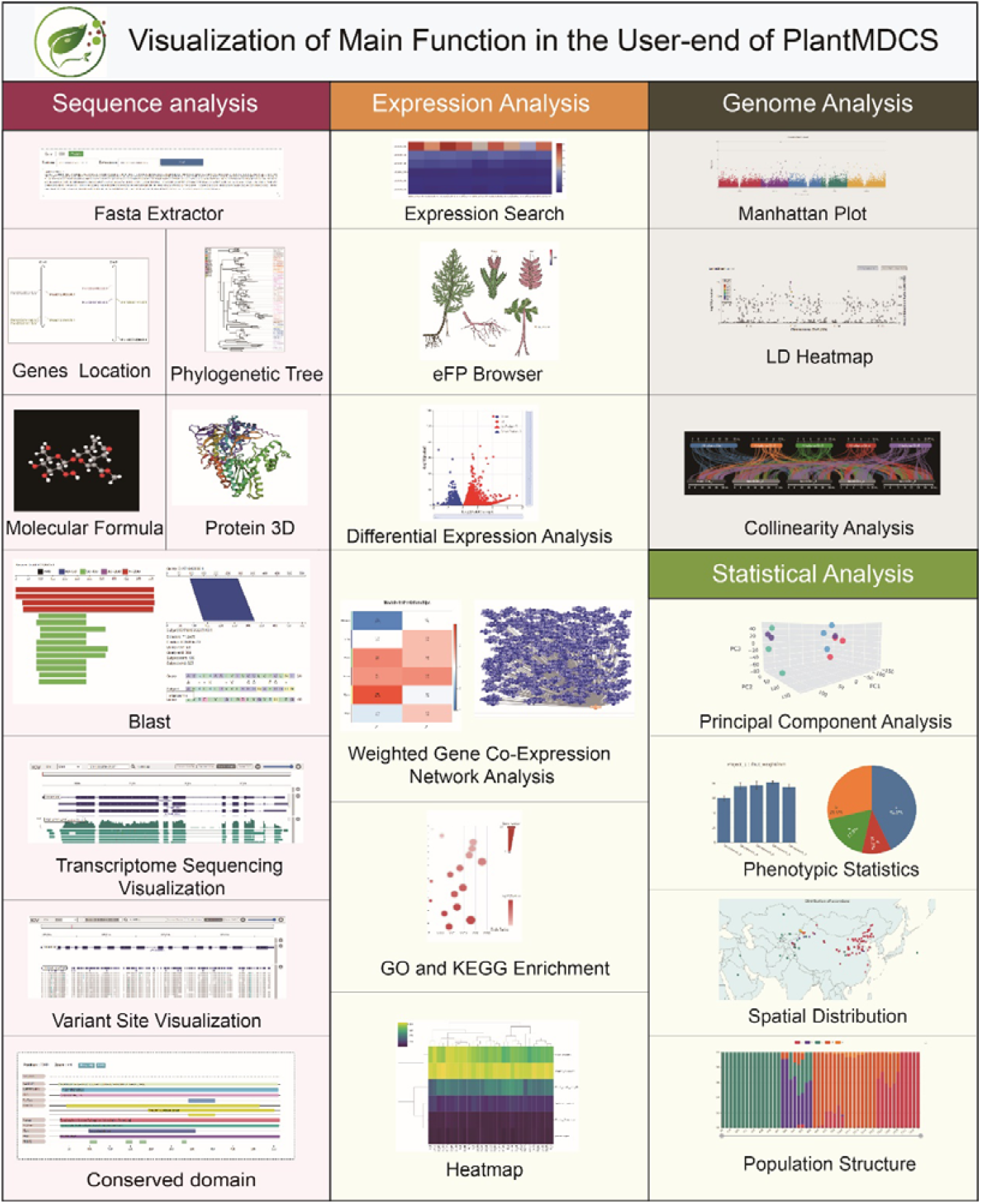
Visualization of main function in the user-end of PlantMDCS. PlantMDCS’s user-end offers multi-omics analysis and diverse visualization effects, with interactive functions for each plot.

### Evaluation of the practical applicability and scalability of PlantMDCS

To rigorously assess the practical applicability and scalability of PlantMDCS, we established locally deployable multi-omics databases for different plant species with genome sizes ranging from compact diploid genomes (e.g., *Oryza sativa*, 363 Mb; *Populus trichocarpa*, 419 Mb) to complex polyploid and repetitive genomes, including *Zea mays* (∼2.0 Gb), *Arachis hypogaea* (° 2.4 Gb), *Saccharum spp*. (up to 10.2 Gb), and hexaploid wheat (*Triticum aestivum*, 13.8 Gb) (Table 2).

**Table 2.**
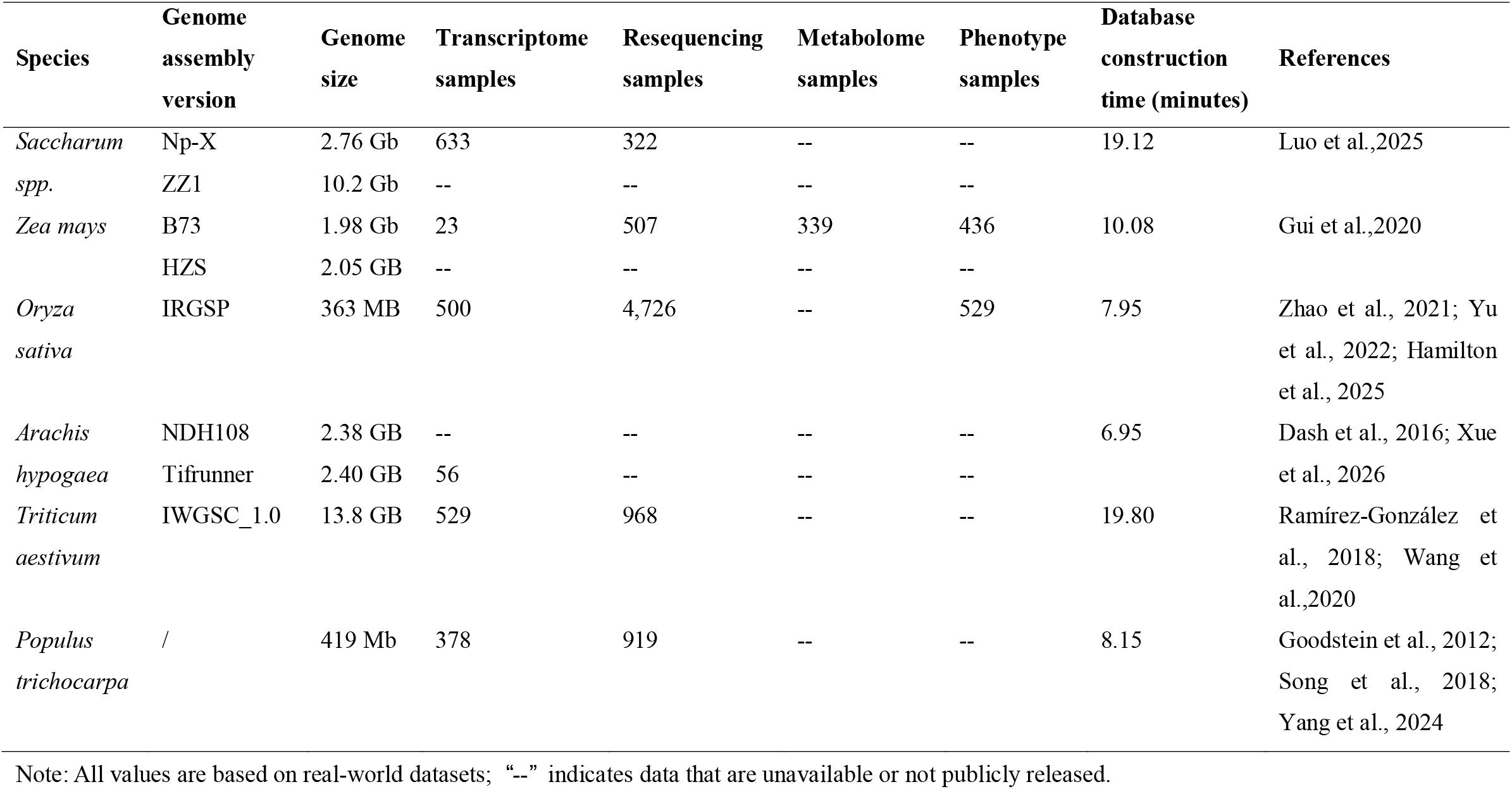
Evaluation of PlantMDCS for local construction of plant 671 multi-omics databases across species.

Across these species, PlantMDCS successfully integrated heterogeneous multi-omics datasets encompassing transcriptomic, population-scale resequencing, metabolomic, and phenotypic data, with dataset sizes and counts varying substantially among species. Notably, complex cases such as rice and maize involved thousands of transcriptome or resequencing datasets and hundreds of phenotypic or metabolomic datasets, reflecting real-world data volumes comparable to those used in established public resources. Despite these differences in genome size and data composition, database construction was completed within minutes for all tested species, demonstrating that PlantMDCS maintains high efficiency across both small and large genomes as well as across varying levels of multi-omics complexity.

Importantly, the observed construction time did not scale linearly with genome size or the number of integrated datasets, indicating that PlantMDCS is not constrained by species-specific genome architecture or project-specific data composition. Instead, the system relies on a genome-centered organizational logic and standardized data ingestion workflows, enabling consistent performance across species with available reference genomes. Together, these results demonstrate that PlantMDCS is applicable to a broad spectrum of plant research scenarios, ranging from model organisms to highly repetitive and polyploid crop genomes, and can support multi-omics data integration at a scale comparable to existing public databases while retaining the advantages of local deployment and rapid construction.

## Discussion

The rapid expansion of plant multi-omics data has fundamentally altered the scale and complexity of biological research, yet the dominant paradigm for data management remains largely unchanged (Luo et al., 2024). Most existing solutions implicitly assume that effective data integration must rely on centralized, server-hosted databases or cloud-based infrastructures, an assumption that introduces persistent risks related to data security, sustainability, and accessibility (Yang et al., 2026). While public databases have undeniably accelerated data sharing and community-wide reuse, they are poorly suited for managing unpublished, proprietary, or continuously evolving datasets generated within individual research groups. Our comparative analysis highlights that even frameworks explicitly designed to support localized deployment, such as e!DAL and Tripal, remain constrained either by limited analytical capability or by substantial technical overhead. These limitations suggest that the challenge is not merely technical but conceptual: current systems prioritize either storage or extensibility, often at the expense of usability, long-term maintainability, and routine analytical integration.

PlantMDCS addresses these limitations by introducing a deliberate shift in analytical logic, from file-oriented data processing to gene-centered, data-driven exploration. Rather than treating multi-omics datasets as loosely connected collections of files requiring repeated preprocessing and manual coordination, PlantMDCS organizes heterogeneous data layers around a genome-centered reference framework that mirrors the biological flow of information from sequence to expression, variation, and phenotype. This design choice has important practical consequences. Once data are ingested, subsequent analyses no longer require users to repeatedly handle raw files or configure pipelines, allowing researchers to focus on biological interpretation rather than technical execution. In this sense, the contribution of PlantMDCS lies not in expanding the number of available analytical functions, but in redefining how multi-omics data are accessed, contextualized, and interrogated. By embedding analytical workflows directly within a persistent data structure, PlantMDCS enables more efficient hypothesis generation and cross-omics reasoning than conventional GUI-based tools that remain fundamentally file-centric.

Equally important is the practical impact of PlantMDCS on long-term usability and sustainability. Our empirical evaluations demonstrate that database construction time is reduced by several orders of magnitude compared with conventional framework-based approaches, primarily due to the elimination of coding requirements, automated data integration, and direct ingestion of large local files. This efficiency gain is not merely a matter of convenience; it directly affects whether multi-omics databases can be realistically constructed, maintained, and reused by small or non-computational research teams. Moreover, by decoupling deployment from ongoing data management and enabling incremental updates through a graphical administrator interface, PlantMDCS substantially mitigates the risks associated with personnel turnover and limited technical support. At the same time, its locally deployable architecture reconciles data security with collaborative accessibility, allowing sensitive datasets to remain under full user control while still supporting controlled internal and remote access. Together, these features position PlantMDCS as a practical and sustainable alternative to both heavyweight public databases and technically demanding framework-based solutions, particularly for plant research groups seeking to balance data ownership, analytical depth, and long-term viability.

## Methods

### Implementation of PlantMDCS

PlantMDCS is implemented using the Python Flask framework combined with the PyQt5 toolkit, with MySQL serving as the relational database management system. The web-based front end is developed using Bootstrap, jQuery, and LayUI to provide a responsive and interactive user experience. The system is designed for local deployment on Windows operating systems and is compatible with mainstream web browsers, including Google Chrome, Mozilla Firefox, Microsoft Edge, and 360 Browser.

### Admin-end implementation for multi-omics database construction

At the administrator end, LayUI is employed to enable graphical file upload through point-and-click interactions. Large data files, including BAM and VCF formats, are supported via chunked uploading and breakpoint resumption using webuploader.js, ensuring robustness during large data transfers. Uploaded data are automatically preprocessed and organized through integrated bioinformatics tools and Python packages. Specifically, samtools (v1.20) (Danecek et al., 2021) and tabix (v0.2.5) (Li, 2011) are used to generate indexes for genome sequence files and to sort and index genome annotation (GFF) files. Gene sequences, coding DNA sequences (CDSs), and protein sequences are extracted from annotated genomes using gffread and Biopython. Sequence homology search databases are constructed using BLAST (v2.15.0) (Camacho et al., 2009).

Structured storage of heterogeneous multi-omics data—including gene expression profiles, protein annotations, transcription factor information, phenotypic measurements, sample metadata, and metabolite structural information—is implemented through MySQL using the pymysql driver, with pandas and numpy facilitating standardized data organization and management. To support system maintenance and task scheduling, real-time monitoring of computational resources (CPU usage, GPU load, and memory consumption) is implemented using psutil and gpustat. In addition, ueditor.js provides a visual interface for homepage configuration and content management at the admin end.

### User-end implementation and core analysis modules

The user-facing interface is designed using Bootstrap and jQuery, enabling users to perform database queries and analyses through graphical interactions without manual file handling. To support concurrent multi-user access and computationally intensive analyses, asynchronous task execution is implemented using Celery and Redis, allowing analysis jobs to be submitted, queued, and retrieved efficiently. The user end comprises two primary functional modules: sequence analysis and multi-omics analysis.

### Sequence analysis module

The sequence analysis module integrates multiple bioinformatics tools and visualization libraries. Biopython is used to extract genomic, CDS, protein, and upstream or downstream promoter sequences. MCScanX (v1.0) (Wang et al., 2012) is employed for synteny analysis between genomes, while BLAST supports nucleotide and protein sequence alignment. Phylogenetic analysis of transcription factor families is automated by combining BLAST, Muscle (v3.8.31) (Edgar, 2004), and FastTree (v2.1.11) (Price et al., 2012), with Biopython (Cock et al., 2009) and ElementTree handling tree format conversion.

Interactive visualization of sequence-related results is implemented using specialized JavaScript libraries: IGV.js (Robinson et al., 2023) for displaying genomic structures, transcriptome BAM files, and resequencing VCF data; D3.js for chromosome localization and BLAST result visualization; Synvisio.js for synteny visualization; Protvista.js for protein domain annotation; 3Dmol.js (Rego and Koes, 2015) and web3Dmol.js (Shi et al., 2017) for three-dimensional protein structure rendering; and PhyloXML.js (Han and Zmasek, 2009) for interactive phylogenetic tree visualization.

### Multi-omics analysis module

The multi-omics analysis module integrates Python- and R-based analytical toolkits to support comprehensive data analysis, including clustering, differential analysis, functional enrichment, time-series analysis, and multi-omics association. Functional enrichment analyses based on Gene Ontology (GO) and Kyoto Encyclopedia of Genes and Genomes (KEGG) pathways are conducted using clusterProfiler. Weighted gene co-expression networks for transcriptomic data are constructed using WGCNA (Langfelder and Horvath, 2008), while Mfuzz (Kumar and Futschik, 2007) is applied for time-series clustering of multi-omics datasets. Differential expression analysis and principal component analysis (PCA) for transcriptomic data are performed using DESeq2 (Love et al., 2014). Phenotypic PCA and clustering analyses are implemented using scikit-learn and seaborn. For population-scale variation analysis, BCFtools (Danecek et al., 2021) is used to extract variant sites, and PLINK (v 0.76) is employed to calculate linkage disequilibrium (LD) between genomic variants.

Analysis outputs are visualized using multiple interactive plotting libraries. D3.js, ECharts.js, and Plotly.js are used to generate bar plots, volcano plots, correlation heatmaps, PCA plots, and Manhattan plots. LocusZoom.js (Boughton et al., 2014) enables visualization of LD patterns within genomic regions, while Heatmaply.js (Galili et al., 2018) produces interactive expression heatmaps. Tabular presentation of raw omics data and analysis results is implemented using DataTables.js.

## Data availability

The executable program, example files, user manual, and instructional videos are available for download from https://github.com/Chen-Chen-888/PlantMDCS. Chinese users can also obtain these files through https://gitee.com/chen-chennnnn/PlantMDCS.

## Funding

This project was supported by Major Scientific and Technological Project of Xinjiang (2024A02006), National Natural Science Foundation of China (grant no.32370243), Science and Technology Projects in Guangzhou (grant no. 2025A04J4383), and Tianchi Talent Project of Xinjiang.

## Author contributions

Y.F., C.C. and J.T. lauched and coordinated the project. C.C. and D.X. developed PlantMDCS software. Y.F., W.L., Y.L., J.S., Y.W., W.Y. and Z.L. collect data and test PlantMDCS functions. C.C., D.X. and Y.F. wrote the manuscript. All authors revised the manuscript.

## Conflict of interest statement

The authors declare no competing interests.

## Notes

### Competing Interest Statement

The authors have declared no competing interest.

